# Emergence of a multidrug-resistant *Salmonella enterica* serovar Amager lineage carrying the *bla*_CTX-M-65_-positive pESI megaplasmid

**DOI:** 10.64898/2026.03.02.708915

**Authors:** Josefina Miranda-Riveros, Daniel Tichy-Navarro, María José Navarrete, Angélica Reyes-Jara, Magaly Toro, Juan A. Ugalde, Andrea I. Moreno-Switt, Alejandro Piña-Iturbe

## Abstract

The spread of extended-spectrum β-lactamase (ESBL)-producing and fluoroquinolone-resistant *Salmonella* pose a global public health challenge in addition to the high burden of infections associated with this foodborne pathogen. In this study we aimed to characterize a multidrug-resistant strain of *Salmonella* serovar Amager isolated from a Chilean river in October 2023. Antimicrobial susceptibility testing revealed a resistance phenotype against multiple antibiotic families, including fluoroquinolones and β-lactams, showing ESBL production. Hybrid genome sequencing allowed the identification of a 311,303 bp plasmid carrying the *aadA1, aph(4)-Ia, aac(3)-IV*a, *floR, sul1, tet*(A), and *bla*_CTX-M-65_ genes, sharing 99.98% sequence identity with the *Salmonella* Infantis pESI-like megaplasmid. In addition, the *qnrB19* gene was found in a ≈2.7 kbp plasmid of widespread distribution. Population structure and temporal phylogenetic analysis at the global scale revealed the emergence of a *Salmonella* Amager lineage from the HC20_35565 cluster, carrying the *Salmonella* Infantis *bla*_CTX-M-65_-positive pESI-like megaplasmid and causing human infections in the United States and the United Kingdom. Our work describes the emergence of a *Salmonella* lineage with resistance against first-line antibiotics used for treating severe infections, underscoring the relevance of environmental surveillance as a means for detecting emergent pathogens and anticipating human infections.

## 1. Introduction

Non-typhoidal *Salmonella* is among the major causes of foodborne illness worldwide. Although typically causing self-limited gastroenteritis, this pathogen can produce severe invasive infections in susceptible groups, such as children, the elderly and immunocompromised people [1], and was responsible for 215,000 estimated deaths only in 2019 [2]. Third-generation cephalosporins and fluoroquinolones are the first-line treatment for severe salmonellosis [1]. However, the increasing antibiotic resistance and the spread of antimicrobial resistance (AMR) genes in *Salmonella* have become major concerns, positioning this foodborne pathogen as a high/critical priority pathogen [3]. In the last three decades, emergent *Salmonella* serovar Infantis has emerged and disseminated worldwide [4,5]. Its success results from the acquisition of ≈ 300 kbp pESI megaplasmid and its variants (pESI-like megaplasmids), which encode multiple factors that favor the virulence, persistence and antibiotic resistance of *Salmonella* Infantis [6,7]. More recently, in the last decade, pESI-like variants carrying the *bla*_CTX-M-65_ gene emerged in the Americas, conferring an extended-spectrum β-lactamase (ESBL)-producing phenotype to the dominant *Salmonella* Infantis subpopulation in the continent [5].

As conjugative elements, the pESI/pESI-like megaplasmids have a potential broad host range among *Salmonella* serovars. Experimental evidence suggests that pESI does not entail a significant fitness cost for its maintenance, and different serovars were found to receive the megaplasmid in conjugations experiments *in vitro* [8,9]. Accordingly, while still uncommon, different reports have documented natural acquisition of pESI megaplasmids by different Salmonella serovars, such as Agona, Alachua, Muenchen, Senftenberg, and Schwarzengrund. The public health and economic impacts caused by the global dissemination of pESI-positive *Salmonella* Infantis underscores the need to carry out surveillance efforts aiming to anticipate the emergence of MDR *Salmonella* serovars driven by pESI or other genomic elements. Here, we present the characterization of an ESBL-producing fluoroquinolone-resistant *Salmonella* Amager strain, isolated from river water in Central Chile, providing evidence of the recent emergence of a *Salmonella* Amarger lineage harboring the *Salmonella* Infantis pESI-like megaplasmid, causing international cases of human infection.

## 2. Materials and methods

### 2.1. Isolation of *Salmonella* Amager 19-MAP-20-4

*Salmonella* Amager 19-MAP-20-4 (antigenic formula: 3,{10}{15}:y:1,2[z45]) was isolated from fresh water sampled from the Mapocho river as part of a surveillance project of *Salmonella* in surface waters (October 5^th^ 2023, Región Metropolitana, Chile) [10,11]. Briefly, 10 L of surface water were collected using a modified Moore swab [12] and a peristaltic pump. The swab was enriched in 200 mL of modified peptone water for 24 h at 37°C and then 100 μL were inoculated in 10 mL of Rappaport-Vassiliadis broth. After 24h of incubation at 42°C, 10 μL were used for isolation in XLT-4 agar using a calibrated bacteriological loop. After 24h at 37°C, three to ten presumptive *Salmonella* colonies were selected for PCR detection of the *invA* gene. Positive *Salmonella* isolates were stored at - 80°C in TSB-glycerol (20% v/v).

### 2.2. Detection of pESI and AMR genes/mutations

Illumina short-read sequencing of isolate 19-MAP-20-4 was available in the public database with the Sequence Read Archive (SRA) accession number SRR30637950. The assembled genome was downloaded from Enterobase [13] (assembly barcode SAL_PD3928AA_AS). MOB-suite v3.1.9 [14] was used for plasmids identification, reconstruction and typing. ABRicate v.1.0.1 was used to assess the presence of the pESI/pESI-like megaplasmid genes using a custom database with the pN55391 megaplasmid genes (GenBank accession NZ_CP016411.1). The pESI-like megaplasmid was considered present if ≥220 genes were found as described before [5]. Predicted antimicrobial resistance genes and mutations were identified with AMRFinderPlus v3.11.26 (database 2023-11-15.1) [15] within Enterobase; and AMRFinderPlus v4.0.23 (database 2025-07-16.1), stand alone, with the “--organism Salmonella” option, for the other HC50_1458 *Salmonella* Amager genomes (see Section 2.5).

### 2.3. Antimicrobial susceptibility testing

The minimum inhibitory concentrations of 34 antibiotic or antibiotic/inhibitor combinations were assessed against *Salmonella* Amager 19-MAP-20-4. Broth microdilution was performed using the ARGNF and CMV5AGNF Sensititre plates (ThermoFisher) following the manufacturer instructions. Result interpretation was carried out according to the Clinical and Laboratory Standards Institute M100-Ed35 document [16].

### 2.4. Whole genome hybrid sequencing and assembly

Genomic DNA (gDNA) was extracted and purified from an overnight culture of *Salmonella* Amager 19-MAP-20-4 using fenol:chloroform:isoamilic alcohol and sodium acetate/propanol precipitation as described before [17]. Then, gDNA was sent to SeqCenter LLC (Pittsburg, PA) for hybrid whole genome sequencing using Illumina and Nanopore platforms. The kits, software and parameters were those reported by SeqCenter

For short-read sequencing, Illumina libraries were constructed using the Illumina DNA Prep kit and sequencing was carried out on an Illumina NovaSeq X Plus platform in a 300 cycle mode for paired-end reads. Base calling, demultiplexing, adapter removal, and initial quality filtering were conducted using bcl-convert v4.2.4.

For Long-read sequencing, libraries were prepared without PCR amplification using the Oxford Nanopore Technologies (ONT) Ligation Sequencing Kit (SQK-NBD114.96), following the manufacturer’s protocols. Nanopore sequencing was conducted using R10.4.1 flow cells in multiplexed, shared-flow-cell configurations. Sequencing runs were configured for the 400 bps mode. The Dorado v0.5.3 basecaller was used under the super-accurate (sup) and 5mC/5hmC (5mC_5hmC) basecalling models. Dorado demultiplex was used under the SQK-NBD114.96 kit settings to generate the BAM file. SAMtools fastq v1.20 was used to generate the FASTQ file from the BAM file. Residual adapter sequences, potentially remaining after basecalling and demultiplexing, were removed using Porechop v0.2.4.

De novo genome assembly was performed using Flye v2.9.2 with the nano-hq model (high-quality ONT reads). Assembly parameters specified a genome size of 6 Mb and initiated assembly using the longest reads sufficient to achieve approximately 50× coverage (--asm-coverage). Resulting assemblies were polished using Illumina short-read data with Pilon v1.24 under default settings. To minimize artifacts associated with low-quality Nanopore reads, long-read contigs exhibiting an average Illumina read depth of ≤15× were excluded from the final assembly. Contigs were assessed for circularization with Circlator v1.5.5 using ONT long reads.

The final assembly was deposited at DDBJ/ENA/GenBank under the accession number JBUDVL000000000 (BioProject PRJNA1417766; BioSample SAMN55014624). A detailed description of the hybrid genome sequencing and assembly methods, along with the corresponding references, is available in the Supplementary Material.

### 2.5. Construction of a time-measured maximum clade credibility tree

The metadata for the 157 Salmonella Amager isolates, along with the results of MLST, cgMLST, SISTR1, SeqSero2, and assembly statistics, were downloaded. To complete missing metadata related to source and year of isolation, searches were conducted using the SRA accession numbers or strain names in Pathogen Detection database (https://www.ncbi.nlm.nih.gov/pathogens/). Of the 157 genomes, those with N50 >100,000 and coverage >30× were retained, resulting in 151 genomes. These were further filtered to remove redundant isolates based on collection year, source niche, country, and HC5, yielding a final set of 107 genomes.

Isolates belonging to the HC50_1450 cluster (n=63) were selected, excluding two lacking information on year of isolation, resulting in 61 isolates in the final selection. The genomes of these 61 isolates were downloaded from Enterobase, and the genome of isolate PNUSAS214304 (assembly barcode SAL_AE7921AA_AS) was selected as the reference based on its high N50 value (571,768 bp) and high coverage (117x). A whole-genome alignment was generated using Snippy v4.6.0, from which recombination regions were identified using Gubbins v3.3 and subsequently removed with snp-sites v2.5.2. A maximum likelihood phylogenetic tree was constructed from the resulting recombination-free core-SNP alignment using IQ-TREE (GTR-F-G4-ASC) and analyzed in TempEst v1.5.3.

To construct the maximum clade credibility (MCC) tree using the selected HC20_1458 genomes, BEAUti and BEAST v1.10.4 were used. The recombination-free core SNP alignment (61 genomes, 1,242 sites) was loaded into BEAUti, and year-of-isolation information was entered. The GTR-G4 nucleotide substitution model was selected (based on the best-fit model identified by Gubbins), with empirically determined base frequencies.

An uncorrelated lognormal molecular clock was selected, and a Bayesian Skyline coalescent model with seven groups and a piecewise-linear skyline was chosen to model population dynamics. The resulting files were examined with Tracer v1.10.7 to assess chain convergence and adequate ESS size. The trees from the independent runs were combined with LogCombiner v1.10.4 and finally used to infer a time-measured maximum clade credibility phylogeny with TreeAnnotator v1.10.4. The resulting tree was visualized in iTOL v7 and additional metadata was incorporated. A detailed description of methods applied for the temporal reconstruction, along with the corresponding references, is available in the Supplementary Material.

### 2.6. Plasmid reconstruction, dereplication and phylogenetic analysis

The Enterobase database was queried for all the *Salmonella* genomes carrying the *bla*_CTX-M-65_ gene (8,022 genomes as of August 29th, 2025). Representative non-redundant genomes (n=1,805) were selected according to the HierCC clustering in Enterobase, selecting one genome per HC10 cluster (genomes linked by ≤10 cgMLST allele differences). MOB-suite v3.1.9 was used for plasmid typing and reconstruction. Plasmids from the AC358 primary cluster (n=1,548) were then clustered with MGE-cluster v1.1.0 with a perplexity value of 100 and a min_cluster value of 5 to create clusters and compute the embeddings.

For the phylogenetic analysis the quality of the reconstructed plasmid was assessed with QUAST v5.3.0 and sequence similarity to the *Salmonella* Infantis reference plasmid pN55391 (accession NZ_CP016411.1) was screened with ABRicate v1.0.1, at the same they were dereplicated with dRep v3.6.2 to choose the best representative genome per cluster producing the set of representatives from which 93 Salmonella Infantis AC358 and 20 non-Infantis AC358 plasmids were selected for core-gene SNP calling with Snippy v4.6.0, using the pESI-like megaplasmid from strain N55391 as the reference. The resulting 33-coreSNP alignment was used to infer a maximum-likelihood tree with IQ-Tree MPI multicore v2.3.2 using ModelFinder Plus for model selection. Node support was assessed using 10,000 ultrafast bootstrap replicates and 1,000 SH-aLRT replicates. The resulting tree was displayed and annotated in iTOL v7 using a circular layout. A detailed description of the methods applied for plasmids analysis, along with the corresponding references, is available in the Supplementary Material.

## 3. Results and Discussion

### 3.1. Characterization of *Salmonella* Amager 19-MAP-20-4 carrying the pESI-like megaplasmid

*Salmonella* Amager 19-MAP-20-4 was isolated from the Mapocho River in October, 2023, and Illumina short-read sequencing was performed. Bioinformatic screening using MOB-suite identified a plasmid from the AC358 primary cluster, the same cluster that groups the *Salmonella* Infantis pESI-like megaplasmids, suggesting the presence of this mobile genetic element. Minimum inhibitory concentration (MIC) assays revealed an MDR profile consistent with the carriage of an ESBL-encoding pESI-like megaplasmid, including resistance to β-lactams (ampicillin, cefotaxime, ceftriaxone), chloramphenicol, ciprofloxacin, tetracycline, and a MIC for the cefotaxime/clavulanic acid combination 64-fold lower than for cefotaxime alone (**Table 1**). Detection of antimicrobial resistance genes and mutations revealed the presence of the genes *aadA1, aph(4)-Ia, aac(3)-IVa, floR, sul1, tet*(A), and *bla*_CTX-M-65_, in accordance with the observed phenotypes. Hybrid Illumina and Nanopore sequencing (accession JBUDVL000000000) enabled the recovery of a 311,303 bp plasmid carrying the *bla*_CTX-M-65_ gene, sharing 99.98% sequence identity (98% coverage) with the pESI-like megaplasmid pN55391 from *Salmonella* Infantis N55391 (**Fig. S1**). Additionally, we detected the *qnrB19* gene, explaining the ciprofloxacin-resistant phenotype, in a 2,623 plasmid (cluster AB042) previously reported in multiple serovars and countries [18].

**Table 1.** Antimicrobial susceptibility profile of *Salmonella* Amager 19-MAP-20-4.

| Antibiotic | MIC Sensititre |  |  |
| --- | --- | --- | --- |
| Amikacin | <= | 8 | NI |
| Amoxicillin/Clavulanic | <= | 8 | S |
| **Ampicillin** |  | **32** | **R** |
| Ampicillin/Sulbactam |  | 16 | I |
| **Azithromycin** |  | **4** | **S** |
| Aztreonam |  | 16 | R |
| Cefepime |  | 4 | SD |
| **Cefotaxime** |  | **32** | **R** |
| **Cefotaxime/Clavulanic acid** | <= | **0.25** | **ESBL** |
| Cefoxitin | <= | 8 | NI |
| Ceftazidime | <= | 2 | S |
| Ceftazidime/Clavulanic |  | 0.25 | NI |
| **Ceftriaxone** |  | **64** | **R** |
| Cefuroxime (sodium) |  | 16 | NI |
| Cephalothin |  | 32 | NI |
| **Chloramphenicol** |  | **16** | **I** |
| **Ciprofloxacin** |  | **1** | **R** |
| Colistin |  | 4 | R |
| Doripenem | <= | 4 | NI |
| Ertapenem | <= | 1 | I |
| Fosfomycin+Glucose6Phosphate | <= | 32 | NI |
| Gentamicin |  | 16 | NI |
| **Imipenem** | <= | **0.5** | **S** |
| **Levofloxacin** |  | **1** | **I** |
| **Meropenem** | <= | **1** | **S** |
| **Minocycline** | <= | **4** | **S** |
| Nalidixic Acid |  | 16 | NI |
| Nitrofurantoin | <= | 32 | NI |
| Piperacillin/Tazobactam | <= | 8 | S |
| Rifampin |  | 8 | NI |
| Sulfisoxazole |  | 256 | NI |
| **Tetracycline** |  | **32** | **R** |
| Tigecycline | <= | 0.5 | NI |
| **Trimethoprim/Sulfamethoxazole** | <= | **2** | **S** |

NI: no interpreted; S: susceptible; I: intermediate; R: Resistant

### 3.2. Global population structure analysis

To place the Chilean *Salmonella* Amager isolate in the global context, we carried out hierarchical clustering of core-genome multilocus sequence typing (cgMLST) allele-profiles from all global *Salmonella* Amager genomes (n=157; Enterobase; July 29^th^, 2025; **Fig. S2** and **Table S1**). These analyses placed the Chilean isolate within cluster HC20_35565 (linked by ≤20 cgMLST alleles), which also included other *bla*_CTX-M-65_–positive genomes from the United States (n=7) and the United Kingdom (n=1), isolated from human infections between 2022 and 2024.

### 3.3. Temporal analysis of the emergence of ESBL-producing *Salmonella* Amager

The maximum clade credibility tree (MCC) of 61 Salmonella Amager isolates from the cluster HC50_1458 was constructed based on 1,242 recombination-free core SNPs using isolate PNUSAS214304 (Enterobase assembly barcode SAL_AE7921AA_AS) as the reference. A time-measured maximum clade credibility tree was generated with all isolates from the HC50_1458 cluster, which encompasses HC20_35565 and closely related isolates (**Fig. 1**). This phylogeny showed pESI-positive *Salmonella* Amager as a monophyletic clade sharing a common ancestor that emerged around 2010.5 (95% HPD 2004.4-2014.8), followed by dissemination across human and environmental niches in different countries and continents. Importantly, we detected the presence of the *qnrB19* gene in four other genomes from the same HC20_35565 cluster, suggesting that this emergent lineage may also be associated with fluoroquinolone-resistance in addition to ESBL production. To investigate the origin of the HC20_35565 *Salmonella* Amager pESI-like megaplasmid, pESI plasmids (cluster AC358) carrying *bla*_CTX-M-65_ were identified from Enterobase assemblies using MOB-suite and were clustered with MGE-cluster (n=1548; August 20^th^, 2025; **Table S2**). The analyses revealed the presence of ESBL-encoding pESI-like megaplasmids among multiple serovars. The pESI-like elements from *Salmonella* Amager clustered very close to each other and within *Salmonella* Infantis-dominated subclusters (**Fig. 2**), in agreement with a scenario in which HC20_35565 *Salmonella* Amager acquired their pESI-like element from a *Salmonella* Infantis strain. Nevertheless, a reference-based phylogenetic reconstruction based on core single-nucleotide polymorphisms (SNPs) of 113 selected pESI megaplasmids revealed a high similarity (0 to 18 SNP differences) among the representative megaplasmids, and no biologically explanatory clusters/clades could be observed (**Fig. S3**).

**Figure 1.**
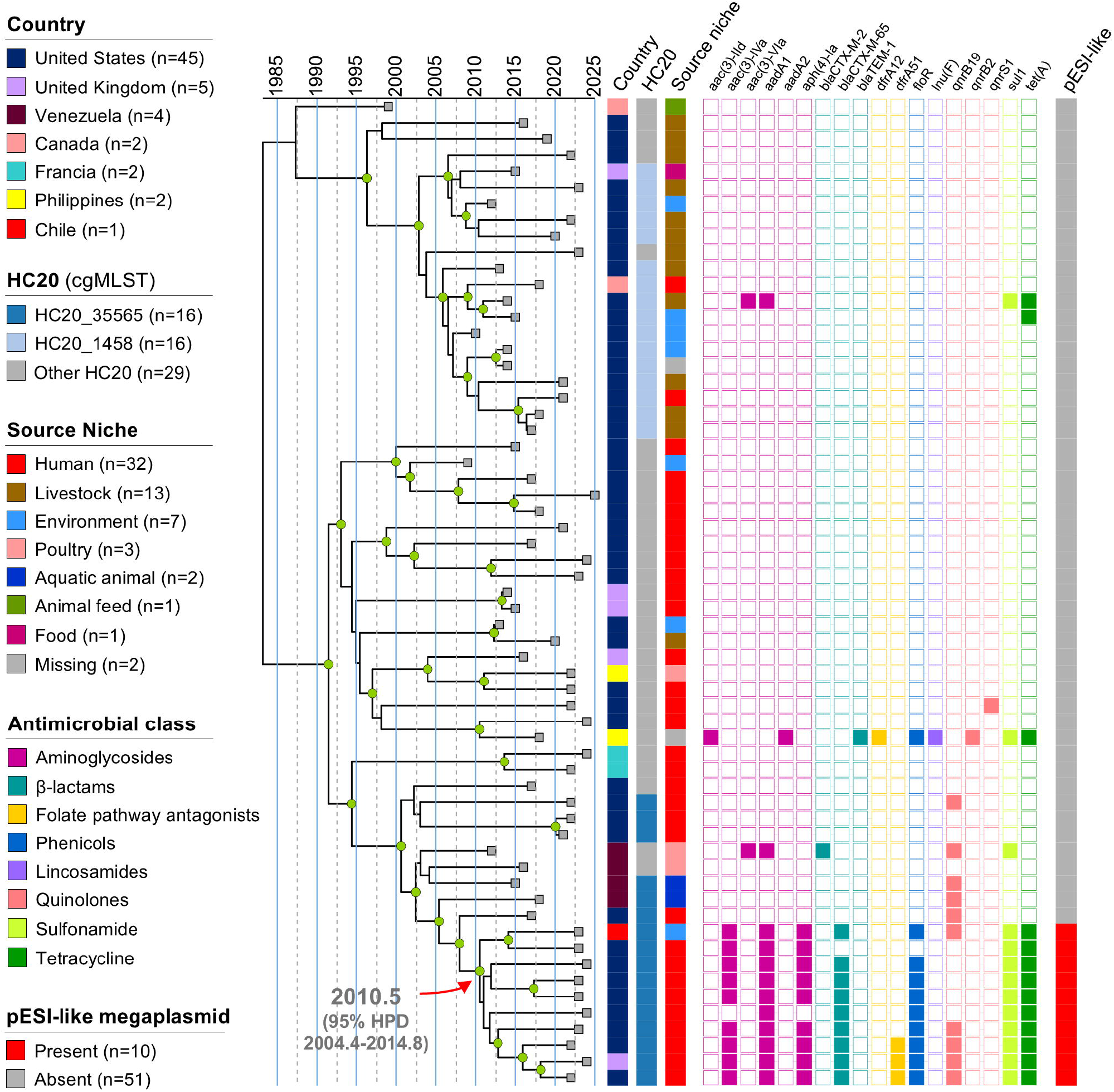
Phylogenomic and temporal analysis of HC50_1458 *Salmonella* Amager. A maximum clade credibility tree (MCC) of 61 *Salmonella* Amager isolates from the cluster HC50_1458 was constructed based on 1,242 recombination-free core SNPs. Internal nodes highlighted with green circles indicate those with posterior probabilities values ≥ 90. Metadata referring to the country of isolation, HC20 cluster, source niche, AMR genes colored by antimicrobial class, and presence of the pESI-like megaplasmids is also shown. The time for the most recent common ancestor for the clade harboring pESI-like megaplasmids is indicated together with the 95% high posterior density (95% HPD) interval.

**Figure 2.**
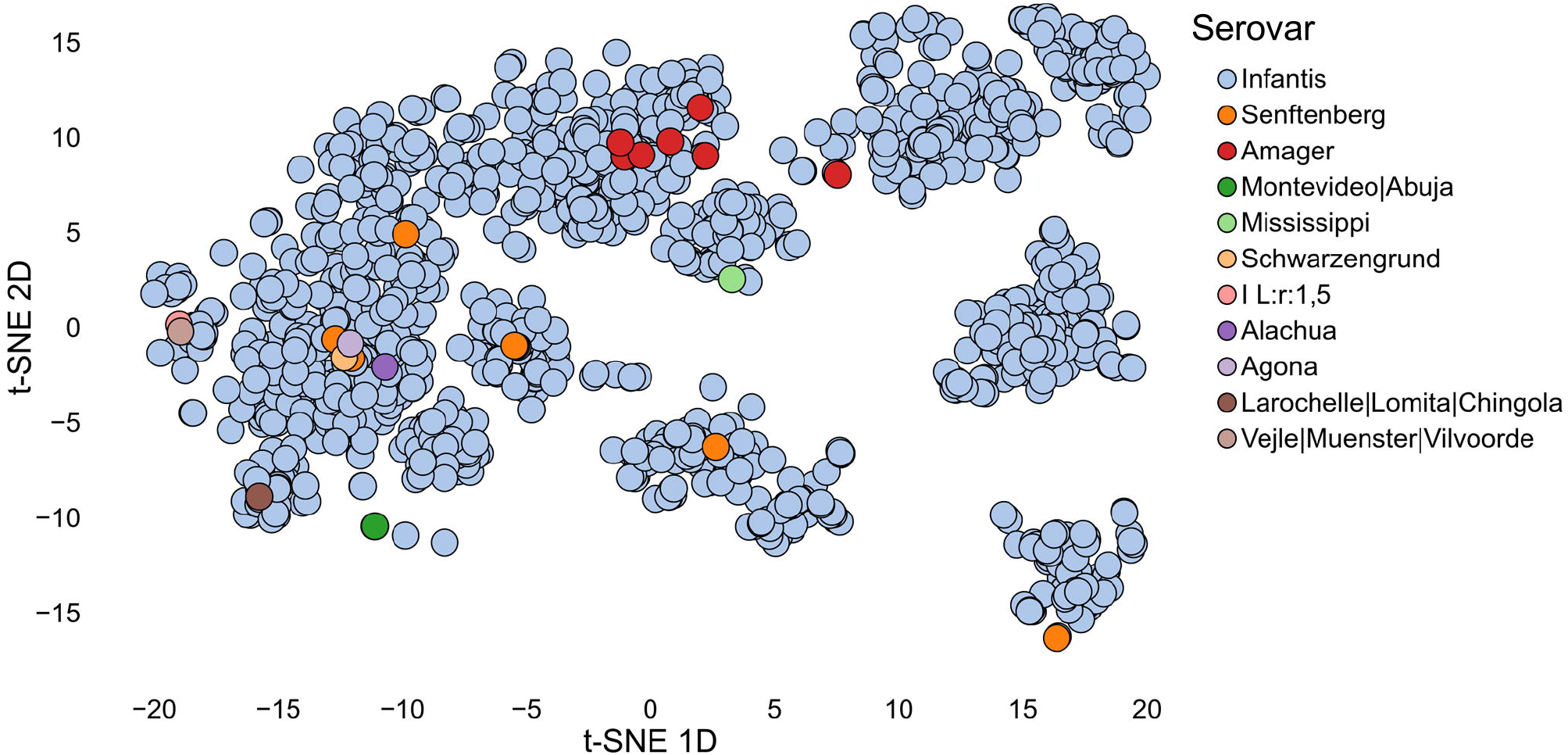
Unitig-based similarity clustering of global *bla*_CTX-M-65_-positive pESI-like megaplasmids. Two-dimensional t-SNE projection of unitig-based profiles for pESI-like megaplasmids (MOB-cluster AC358) reconstructed from *Salmonella enterica* assemblies carrying *bla*_CTX-M-65_ identified in EnteroBase. Points are colored by serovar as reported by SISTR1 in Enterobase (color key shown in the inset legend).

## 4. Conclusion

The emergence of non-Infantis pESI-carrying *Salmonella* has been previously reported. In 2022, MDR *Salmonella* Munchen (lacking the *bla*_CTX-M-65_ gene) was described in Israel, associated with horizontal acquisition of a pESI-like megaplasmid [9]. In the same year, pESI-like elements were identified in the serovars Agona, Munchen, Schwarzengrund, and Senftenberg [19]. More recently, in 2024, pESI-mediated ESBL-mediated cephalosporin resistance was reported in the United States in serovars Senftenberg, and Alachua, further demonstrating the plasmid capacity for horizontal transfer among diverse serovars [20]. The spread of the *bla*_CTX-M-65_ gene to other *Salmonella* serovars is a major concern and, as observed with HC20_35565 *Salmonella* Amager, its co-occurrence with fluoroquinolone-resistance determinants further limits critical first-line treatment options for severe salmonellosis. The isolation of ESBL-producing pESI-positive *Salmonella* Amager in different countries (and continents) highlights the global threat posed by *bla*_CTX-M-65_-carrying pESI-like megaplasmids. Our work also underscores the relevance of environmental surveillance as a means for detecting emergent pathogens and for anticipating human infections. Collectively, current data should alert AMR and *Salmonella* surveillance systems, health-care professionals, and stakeholders to the emergence of ESBL production across multiple serovars.

## Funding

Powered@NLHPC: This research was partially supported by the supercomputing infrastructure of the NLHPC (CCSS210001).

Agencia Nacional de Investigación y Desarrollo de Chile -ANID-Fondecyt Postdoctorado Folio 3230796.

This research is partially supported by the FDA of the U.S. Department of Health and Human Services (HHS) as part of a financial assistance award U01FDU001418 in the scope of the Cooperative Agreement to support the Joint Institute for Food Safety and Applied Nutrition (JIFSAN).

## Acknowledgments

The authors of this work were supported by Agencia Nacional de Investigación y Desarrollo de Chile -ANID-through grants Fondecyt Postdoctorado Folio 3230796, Fondecyt Regular Folio 1231082, FONDEF IDeA I+D Folio ID24I10291, and scholarships Beca de Doctorado Nacional Folio 21241909 and Folio 21251661.

## Ethical Approval

Not required.

## Data availability

All genome sequences are publicly available. Accession numbers and associated metadata are listed in the Supplemental Tables.

